# A Molecular Classification of Human Mesenchymal Stromal Cells

**DOI:** 10.1101/024414

**Authors:** Florian Rohart, Elizabeth Mason, Nicholas Matigian, Rowland Mosbergen, Othmar Korn, Tyrone Chen, Suzanne Butcher, Jatin Patel, Kerry Atkinson, Kiarash Khosrotehrani, Nicholas M Fisk, Kim-Anh Lê Cao, Christine A Wells

## Abstract

Mesenchymal stromal cells (MSC) are widely used, isolated from a variety of tissues and increasingly adopted for cell therapy, but the identity of these cells is poorly defined and commonalities between MSC from different tissues sources is controversial. Here we undertook a comprehensive review of all public MSC expression studies to assess whether cells derived from different sources shared any common molecular attributes. In doing so, we discovered an over-arching transcriptional phenotype shared by a wide variety of MSC, freshly isolated or cultured cells, and under a variety of growth conditions. We developed a modified variable selection protocol that included cross platform normalisation, and assessment of the selected gene stability and informativeness. A 16-gene signature classified MSC with >97% accuracy, discriminating these from fibroblasts, other adult stem/progenitor cell types and differentiated cells. The genes form part of a protein-interaction network, and mutations in more than 65% of this network were associated with Mendelian disorders of skeletal growth or metabolism. The signature and accompanying datasets are provided as a community resource at www.stemformatics.org resource, and the method is available from the CRAN repository.

## Introduction

Adult tissues maintain the capacity to be replenished as part of the normal processes of homeostasis and repair. The adult stem cell hypothesis proposes that multipotent cells resident in tissues are the source of this cellular renewal, and expand in response to tissue injury. MSC were first isolated from bone marrow, where these occupy an important stem cell niche required for reconstitution of bone and the stromal compartments of marrow, and also play a supportive role in haematopoiesis (*1*, *2*). Subsequently adult stromal progenitors have been isolated and cultured from most organs including placenta, heart, adipose tissue and kidneys although the identity of these cells remains controversial (reviewed by (*3*, *4*)). Specifically the question of how similar cells isolated outside the bone marrow niche are is unresolved, nor whether these could be considered bona fide MSC, or indeed challengingly, whether MSC isolated from different tissues share any phenotypic or molecular characteristics at all (*3*). In this light various cells described as MSC (whether by name or attribution) have been reported as having quite different self-renewal capacity, immunomodulatory properties or propensity to differentiate *in vivo* (*5*). It has been variously argued that MSC isolated from most stromal tissues are derived from perivascular progenitors (*6*), or recruited from the bone marrow to distal tissue sites (*7*), or that resident stromal progenitors from different tissues must have tissue-restricted phenotypes.

The question of ontogeny aside, there is little consensus on whether MSC from differing tissue sources share common functional attributes. Most human studies have been conducted on very small numbers of donors, so it is difficult to dissect donor-donor heterogeneity from source heterogeneity. Donor-donor variation is a major contributor to differences in MSC growth and differentiation capacity, and clonal variation is evident even when derived from the same bone marrow (*8*, *9*). Consequently there is little agreement in the literature on definitive molecular or cellular phenotypes of human cultured MSC, whether from bone marrow or other sources.

The lack of definitive markers for human MSC is a major barrier to understanding genuine similarities, or resolving differences between various cell sources or subsets. Modern molecular classification tools are needed for the characterisation of MSC ex vivo and in vivo. Here we describe a sophisticated integrative transcriptome analysis of public MSC datasets to assess how similar these cells are, and describe the major classes of MSC captured in the literature to date.

## Results

### A 16-gene MSC signature defines an overarching MSC phenotype

To address whether MSC shared a common molecular phenotype that could distinguish them from other stromal or progenitor cells, we systematically reviewed all of the publicly available transcriptome data for presumed MSC, identifying 120 potential datasets that were derived from a wide variety of tissues and age groups, but 35/120 datasets failed our QC criteria for data quality and were excluded from the study. We assessed the accompanying phenotypic data of the remaining 85 datasets carefully (Supplementary Tables S1-S5) for immunophenotype and ex vivo differentiation, as determined by the International Society for Cellular Therapy (*10*). A ‘gold standard’ sample set was assembled, consisting of 125 MSC samples from 16 independently derived datasets. The ‘Gold standard’ MSC were primarily derived from bone marrow, but also included a variety of adult, neonatal and fetal stromal sources, and these were compared to 510 non-MSC samples from primary human tissues, including cultured fibroblasts, haematopoietic cells and pluripotent stem cell lines (Supplementary Table S1, S2).

We derived a novel cross-study framework to test whether we could find similarities between the MSC in our training set despite tissue, platform or laboratory differences. Our approach, described in Figure 1A, included a cross-platform normalisation step (*11*), and a modified multivariate discriminant analysis that included steps to evaluate the stability of gene selection when datasets were subsampled as well as steps to evaluate the informativeness of the variables that contributed to each component (Figure 1B, Supplementary Figure S1).

**Figure 1.**
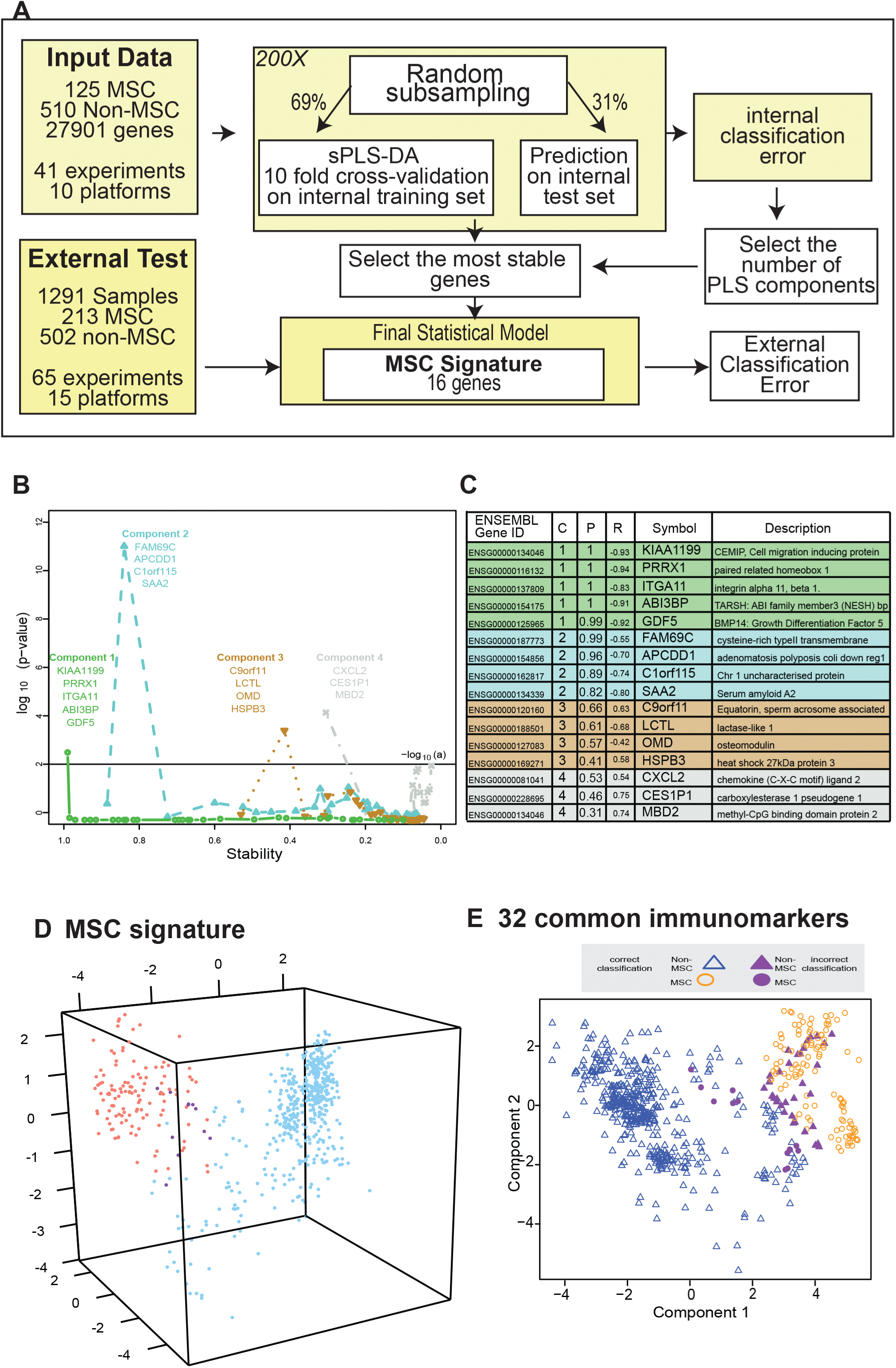
Identifying the MSC signature. A) Workflow summarizing our improved implementation of the sPLS-DA and derivation of a stable 16 gene MSC classifier; B) Choosing the most informative minimal gene set on each component by testing the benefit of including more genes. Each dot is a gene set, ordered along the x-axis by decreasing stability (frequency of selection). The y-axis represents the -log_10_(P-value) of a one tailed t-test at level a=1% indicating the improvement in classification accuracy; Component 1 genes are indicated in green, component 2 genes in blue, component 3 genes in brown and component 4 genes in grey. C) The 16-gene MSC signature colour coded to the component that it contributes to (as per 1B). Gene ID is given as HUGO symbol and ENSEMBL gene ID; ‘C’ is component; ‘P’ is probability of selection (indicating stability); ‘R’ is correlation of gene to component. D) Sample clustering and classification accuracy of the training set (635 samples), clustering shown along the first three components. MSC samples are shown in red, non-MSC in blue, and misclassified samples in purple. E) A panel of 32 commonly used MSC immunomarkers used to cluster the 635-curated samples in the training set. Correctly classified MSC are red circles, correctly classified non-MSC are blue triangles, incorrectly classified samples are purple.

This identified 16 genes (Figure 1C) that collectively formed a ‘signature’, which across 4 components grouped bone-marrow derived MSC with MSC from other sources, and provided a high degree of discrimination between MSC and non-MSC cell types (Figure 1D, Supplementary Figure S1). The accuracy of the signature (Table 1) was 97.8%, calculated as the percentage of correctly classified samples in 200 subsamplings of an internal test set.

**Table 1:** MSC Signature improves the classification accuracy of MSC compared to a panel of 32 commonly used MSC markers.

|  | CD45,<br>CD73,<br>CD105 | 32 common<br>MSC<br>markers | sPLS-DA prior<br>to stable gene<br>selection | The 16-gene MSC<br>signature |
| --- | --- | --- | --- | --- |
| **Overall accuracy**<br>(% of 635 samples) | 87.86 | 92.33 | 97.71 | 97.85 |
| **MSC misclassified**<br>(% of 125 samples) | 14.40 | 11.10 | 3.04 | 4.31 |
| **Non-MSC misclassified**<br>(% of 510 samples) | 11.60 | 6.82 | 2.11 | 1.61 |

Column 1 provides the comparison of the classification accuracy of the 635 training samples using (Column 2) the 3 markers used as the minimal immunophenotype of the MSC training samples; (Column 3) a panel of 32 commonly used immune-markers in the MSC literature; (Column 4) using the unrefined sPLS-DA output; or (Column 5) with our final signature of 16 genes. Performance of each gene group was assessed using 200 random subsamplings of the training set. The internal classification error rate was calculated from a PLS-DA with 2 components (known immune-markers), or was an output of our statistical model with genes selected in an unbiased manner (cf Figure 1A).

### The signature is more informative than a literature-derived set of MSC markers

The International Society for Cellular Therapy (*10*) had identified a set of commonly used markers to immunophenotype MSC (Supplementary Table S6). These markers did cluster MSC samples from the training set together on a MDS plot (Figure 1E), but 6.7% (34/510) of non-MSC samples also clustered with this group, the majority of which (73.5%) were fibroblasts. A further 10.4% (13/125) of MSC samples were misclassified as non-MSC. The proportions were similar over 200 subsamplings of an internal test set (Table 1). This high misclassification rate is consistent with a large body of literature documenting the ambiguity of these markers, which are shared with stromal fibroblasts, endothelial progenitors and hematopoietic cells. The variable expression of these markers (Supplementary Figure S2) may also explain the variability of marker use reported by the wider MSC research community.

### The MSC signature genes are essential to healthy mesenchymal development and function

To assess possible functional relationships between MSC signature genes, we used a curated set of protein-protein interactions from the BioGrid database using the genes selected from component 1 that showed a high discriminating power between MSC and non-MSC. These formed a network of 43 interacting proteins (Figure 2A). A high proportion of these (30/43) are represented in Mendelian disorders of mesenchymal development by virtue of their mutation spectrum in facial or musculo-skeletal dysmorphologies in man, or evidence of mesodermal defects in KO mouse models (Supplementary Table S7). These included the paired-related homeobox-1 (PRRX1), a transcription factor important for early embryonic skeletal and facial development, and with a *de novo* mutation spectrum in the embryonic dysmorphology syndrome Agnathia-otocephaly (*12*). Likewise, mutations in bone morphogenetic protein 14 (BMP14/GDF5) lead to developmental abnormalities in chondrogenesis and skeletal bone (*13*). Mutations in DDR2 cause limb defects, including spondylo-epiphyseal-metaphyseal dysplasia (*14*) and mice over-expressing DDR2 have increased body size and atypical body fat (*15*). Polymorphisms in ABI3BP are associated with increased risk of osteochondropathy (*16*).

**Figure 2:**
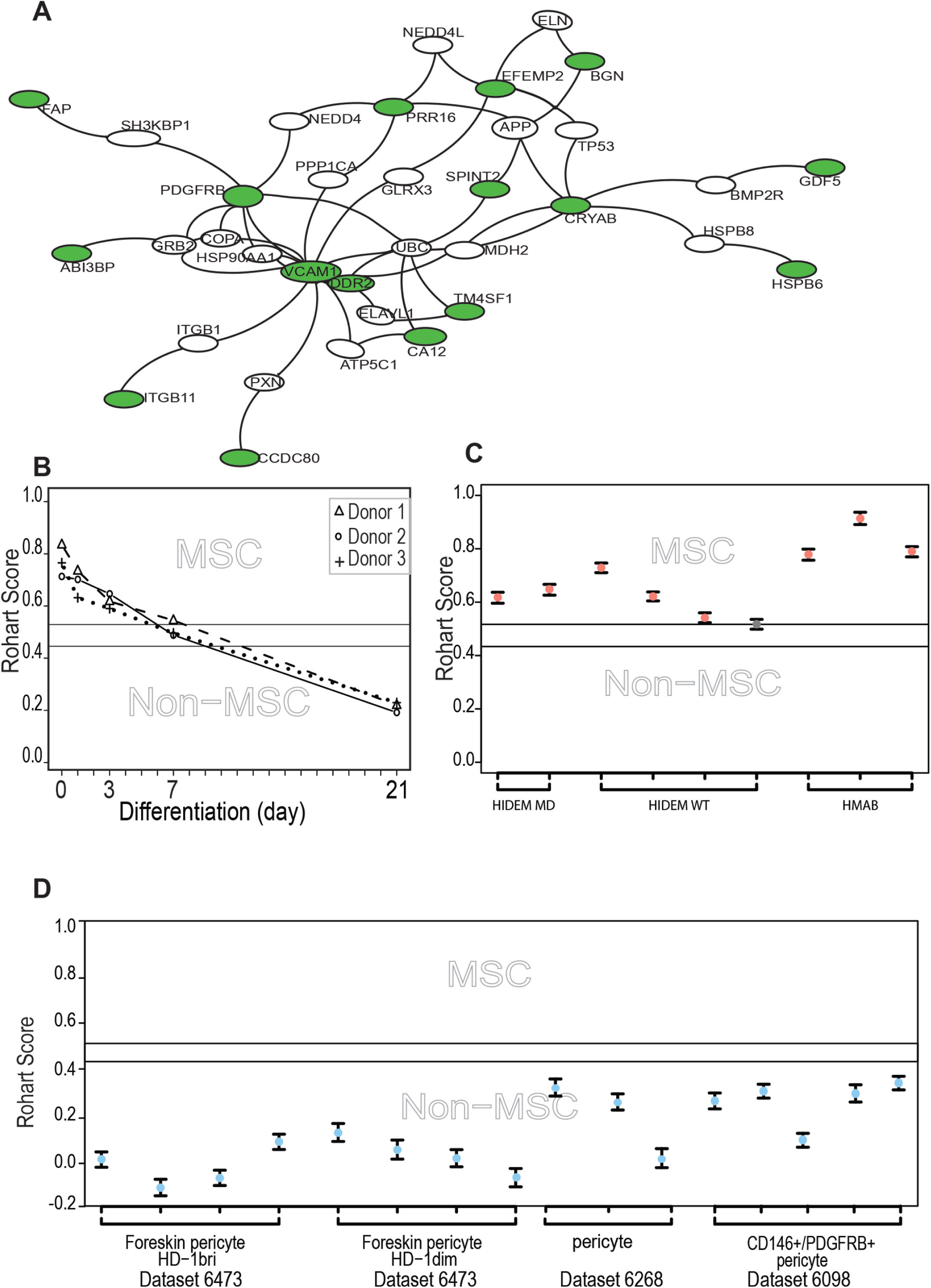
The MSC signature forms part of a network of extracellular proteins and discriminates between differentiating or related adult stem cell types. A) An extended protein-protein network diagram of the Rohart MSC signature genes demonstrating a role for VCAM1 and PDGFRB as part of a functionally interconnected set of glycoproteins, integrins, growth factors and extracellular matrix proteins. Green nodes are seed network members from component 1 of our statistical model, white nodes are inferred network members, and edges are protein-protein interactions. B) Classification of bone marrow MSC over a time course of differentiation to cartilage; y-axis gives the MSC score, x-axis orders the samples from each experimental series. Three differentiation series from three donors are shown. The uncertainty region stands between the MSC and non-MSC prediction regions. C) Classification of perivascular-derived stem cells from skeletal muscle mesangioblasts (HMAB), or iPSC-derived mesangioblasts (HIDEM) from donors with muscular dystrophy (MD) or healthy donors (WT). Error bars around each prediction score represent the CI boundaries. A sample is classified as ‘unsure’ (indicated in grey) if its prediction score or its CI overlapped the uncertainty region. D) Classification of pericytes derived from three distinct datasets: from Left-Right neonatal foreskin (Antigen HD-1 dim or bright); placental pericytes; perivascular endometrial stem cells (CD146+/PDGRFB+). Stemformatics dataset identifiers provided for each experimental series. Error bars around each prediction score represent the CI boundaries.

The extended network included several previously described MSC immune-markers. ITGA11 for example was a member of the core signature, and although it is not a commonly adopted MSC marker, ITGA11 has been used to prospectively enrich MSC from bone marrow with enhanced colony forming capacity (*17*), and independently shown to be enriched more than 3 fold at protein level in bone marrow MSC compared to dermal fibroblasts or perivascular cells (*18*). Other members of our network that have been previously described in human or mouse MSC biology, and validated at the protein level include PDGFRβ (*19*), SPINT2 (*20*), CCDC80 (*21*), FAP (*22*), biglycan (*18*), TM4SF1 (*23*) and VCAM1. SPINT2 is a serine protease inhibitor whose activity is required in bone-marrow MSC, and its loss alters hematopoietic stem cell function in myelo-dysplastic disorders (*20*). In mouse, CCDC80 is also necessary for reconstitution of bone marrow and support of haematopoiesis (*21*). VCAM1 together with STRO-1 has been used for the prospective isolation of human bone marrow MSC (*24*). VCAM1 is an adhesion molecule that is induced by inflammatory stimuli to regulate leukocyte adhesion to the endothelium (*25*); however in cardiac precursors its expression demarcates commitment to mesenchymal rather than endothelial lineages (*26*).

The network included a high proportion of extracellular proteins (54%) with demonstrated roles in the modification of extracellular matrix proteins including proteoglycans, as well as regulators of growth factor and cytokine signalling. This included the cell migration inducing protein (KIAA1199/ CEMIP), which is secreted in its mature form. It regulates Wnt and TGFβ3 signalling by depolarising hyaluronan, and may alter trafficking of cytokines and growth factors to the extracellular milieu (*27*). DDR2 is a receptor tyrosine kinase that interacts directly with collagens. It stabilises the transcription factor SNAIL, and has been implicated in epithelial-mesenchyme transitions in epithelial cancers (*28*). CCDC80 binds syndecan-heparin sulphate containing proteoglycans, has been shown to inhibit WNT/beta-catenin signalling and has a regulatory role in adipogenesis (*29*, *30*). SRPX2 is a secreted chondroitin sulfate proteoglycan involved in endothelial cell migration, tissue remodelling and vascular sprouting (*31*). The chaperonins HSPB5/CRYAB and HSPB6 stabilise protein complexes, and may assist in delivery of growth factor complexes where these are present in high concentrations. In transplantation paradigms it is likely that the therapeutic benefit derived from MSC is via local immunomodulatory, anti-inflammatory, and/or trophic effects during the acute phase of cell therapy. The network of genes identified here as enriched in MSC suggests an over-arching role for these cells in modifying the extracellular environment, functions important in development as well as in homeostatic regulation of adult tissues.

### MSC differentiation, dedifferentiation and the MSC signature

The majority of public microarray datasets available to us had limited phenotypic data available, so these were not used to derive our MSC signature. Nevertheless we annotated each of these samples as *presumptive* MSC (213 samples) or *presumptive* non-MSC (499 samples) based on their origin and use in the source publication (Supplementary Table S4). Where MSC were profiled during *in vitro* lineage differentiation, we assigned the samples taken at intermediate time points to an ‘unknown’ category (579 samples) prior to testing these with the signature.

Despite the lack of phenotypic information associated with these datasets, the agreement between publication status and our classification was high, with only 9 presumptive MSC scored as non-MSC (4.2%). Five percent of the presumptive non-MSC (27/499) were misclassified by the signature as MSC, and around half of these (>13) were neonatal or fetal dermal fibroblasts (Supplementary Table S4, S5), although these formed the cluster with the greatest distance from the bone-marrow MSC. Fibroblasts from other sources were not classified as MSC, furthermore, the signature could discriminate between MSC and differentiating cultures. Figure 2B demonstrates loss of the MSC score during chondrogenic differentiation with the addition of TGFβ3 (Dataset 6119 (*32*)) and this pattern was recapitulated for cells differentiating to mineralising bone or to adipose-like cells or when undergoing reprogramming of an adipose-tissue derived iPSC (Supplementary Figure S3).

### Comparison of MSC and adult stem/progenitor cell types

Some MSC subsets are likely to be derived from a perivascular progenitor. In our hands, primary skeletal-muscle mesoangioblasts (Dataset 6265 (*33*), defined as alkaline-phosphatase^+^ CD146^+^ CD31/Epcam^-^ CD56/Ncam^-^ with demonstrated skeletal muscle differentiation, and which are thought to be a subset of perivascular cells in skeletal and smooth muscle were classified as MSC (Figure 2C). In contrast, the majority of cells derived from a perivascular location (and confirmed as such with tissue imaging) were not classified as MSC (Figure 2D). The limbal cell niche hosts both limbal epithelial and stromal progenitors (*34*), and the stromal progenitors are also classified as MSC by our tool (Dataset 6450).

### Transcription factor expression drives tissue clustering of MSC but is confounded by gender and MHC-1 haplotype

The capacity to group MSC-like cells is consistent with the general assumption that MSC from different tissue share some common molecular properties, but is not otherwise able to assess functional differences between cells isolated from different sources.

When we examined the clustering of MSC by the 16-gene signature (Figure 3), we observed that the majority of bone marrow MSC clustered together, and these were more similar to fetal blood or fetal liver derived cells than MSC isolated from adipose tissue, placenta or skin. We therefore examined more broadly the genes that were significantly different between bone marrow MSC and other cell types at the whole transcriptome level. This analysis confirmed the observed clustering of bone marrow derived MSC, distinguished by differential expression of 425 genes (adjusted P< 0.01, Supplementary Table S8). Gene ontology analysis revealed higher expression of transcription factors associated with skeleto-muscle morphogenesis in BM-MSC (Figure 3C), indeed 25% of the top 100 differentially expressed genes were transcription factors highly expressed in a subgroup of bone-marrow MSC, with known roles in embryonic patterning and organogenesis.

**Figure 3.**
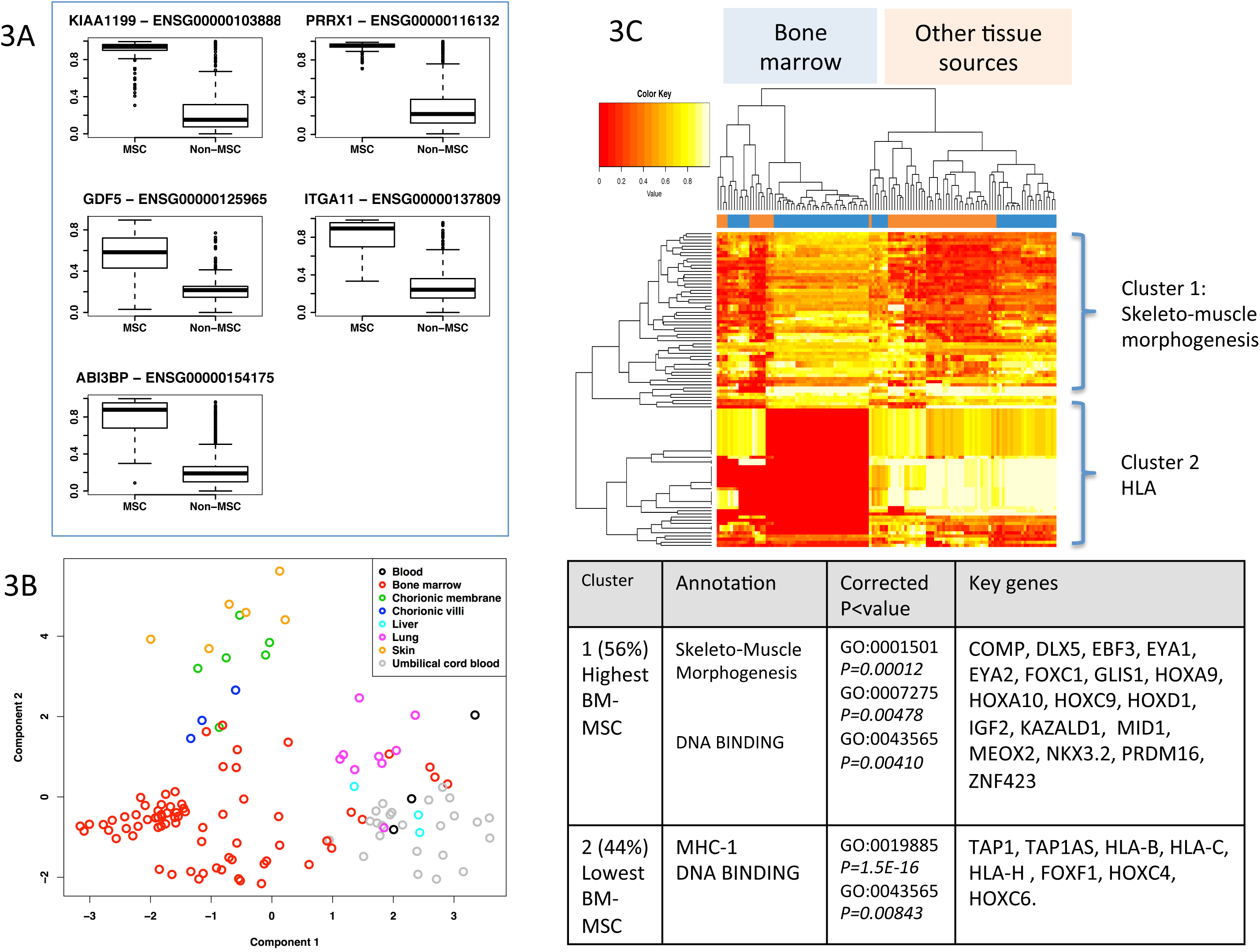
An over-arching MSC phenotype encapsulates distinct subtypes. A) Box and Whisker plots showing average expression of the component 1 genes making up the MSC signature. B) PLS-DA of the MSC samples in the training data set, coloured by tissue of origin. C) Heat map clustering well-annotated MSC samples (columns) with the top 100 differentially expressed genes (rows). Sample tree is coloured Bone-marrow (blue) vs any other tissue (orange). Heat map coloured lowest expression (red) to highest expression (white). GO terms enriched in the heat map clusters and the key genes driving the enrichment terms are provided in a table below the heat map.

Most of the previous reports assessing tissue-dependant differences in gene expression, immunomodulation or differentiation capacity use small numbers of donors and the lack of concordance in the literature may indicate that donor heterogeneity is a major confounder in these prior analyses. The genes that were most differentially expressed between the different MSC sources in our combined analysis were MHC class I genes, and these accounted for >40% of the top 100 differentially expressed genes in the bone-marrow comparisons (Supplementary Table S8 and S9), which suggests donor remains the largest contributor to variation. The HLA isotypes were generally, but not exclusively, expressed at lower levels in bone marrow MSC. Estrogen and progesterone receptors, and a network of associated target genes were also significantly different between tissue sources (Supplementary Table S9), and this may reflect a gender bias in tissue sampling; although the gender of the donors was not available for a majority of samples, some tissues (such as decidual sources) will be entirely female in origin. Further molecular sub-classifications of MSC will therefore much larger studies that address specific clinical or differentiation properties of the cells, and must also consider ascertainment biases that may introduce confounding variables such as HLA subtypes or gender.

All together we curated more than 120 MSC-related gene expression datasets in the www.stemformatics.org resource (*35*); the datasets can be queried here using key word, dataset ID or author, together with an implementation of the Rohart MSC test. We noted that freshly isolated bone-marrow MSC profiled in 3D cultures (datasets 6416 (*36*) and 6544 (*37*)), or over several passages (dataset 6254 (*38*) and dataset 6345 (*39*)), or derived from donors of different ages (gestational - dataset 6063 (*40*) and the elderly - dataset 6216 (*41*)) retained the MSC signature. The signature was robust for cells cultured in the human serum, platelet factors or FBS (dataset 6393 (*42*)), or in cells supplemented with TGFb1 (dataset 6345 (*39*), suggesting that the phenotype defined by this signature is independent of the method of isolation or culture. These data are consistent with a shared molecular phenotype that can classify a wide range of freshly isolated and cultured MSC, but one that is rapidly lost under differentiating conditions.

## Discussion

Our study set out to test the commonly held assumption that MSC from different sources share common and specific properties, as is increasingly challenged in the literature (*3*). In doing so we identified a specific gene signature that is shared by a wide-variety of MSC, but not common non-MSC cell types. We have implemented the resulting classification tool as a simple online test that will be useful in standardisation or improvement of current bulk isolation methods. The “Rohart test” MSC classification tool is available in the Stemformatics.org platform, together with all the primary data used in derivation of the signature. Details on submitting proprietary data to the Rohart test are available on the stemformatics.org site.

Our approach highlights the potential robustness of biological signatures when combining data from many different sources, where experimental variables such as platform or batch can be reduced (Figure S4). The methods we used for derivation of a common MSC classifier could be applied to the meta-analysis of any cell subset or phenotype where sufficient samples can be drawn from public expression databases. The bulk of the expression data available to us was generated on microarray platforms, and this brings some caveats. VCAM1, CD73 and CD105 were part of the extended PPI network but were not included in our core signature. This may be a consequence of cross-platform meta-analysis, as not all MSC genes will be represented with a quality probe on all of the array platforms, or may reflect a lack of specificity when MSC are compared to a wide atlas of other cell types, or may reflect a high variability in mRNA expression from donor to donor or across tissue sources. Nevertheless the inclusion of these known MSC markers in our extended network, and demonstration of the high connectivity between VCAM1 and our core signature, together with the high level of genetic perturbation impacting on mesenchymal tissues in this network, provided us with confidence in the biological relevance of our core members.

The question of what is an MSC, and whether these are a distinct stem cell population recruited from the bone marrow, as suggested by mouse studies of fetomaternal microchimerism (*43*) or from perivasculature, as suggested by immunotagging of MSC-like cells from perivascular regions in human tissues (*6*), or are resident progenitor populations specific to each organ cannot be resolved in the current study. However a particular strength of this signature was the sub-grouping MSC into distinct clusters, including a predominantly bone-marrow cluster. The clustering of MSC, perivascular cells and fibroblasts as distinct cell types is consistent with the idea that these may represent a continuum along a stromal stem cell hierarchy.

The signature itself is obviously dependent on the quality of the MSC used in the training set. In this light, we curated 120 public datasets; 35 failed on QC and 37 lacked sufficient phenotypic annotation for MSC, leaving 125 MSC from 16 independently derived datasets as ‘gold standard’ MSC. As rare adult stem/progenitor cell types were under-represented in the current test or training datasets, we anticipate that functional classification of MSC subtypes will improve as newer sampling methods provide the means to identify and replicate these cells. We expect that further refinements in the isolation or culture of purer MSC or more precisely defined functional subsets will also result in further refinements of this signature.

In summary, we set out to systematically review the current state of play in MSC biology using a meta-analysis of transcriptome studies, and in doing so were able robustly to identify a general MSC phenotype that could distinguish MSC from other cell types. The resulting signature could also identify points of transition as MSC underwent differentiation or reprogramming studies. Furthermore, we demonstrated that, at least at a gene expression level, our *de novo* derived signature outperformed the classification accuracy of the combined set of traditional MSC cell surface markers. While a signature approach such as ours is not able to resolve the ontogeny or in vivo function of MSC, it does provide a tool for better benchmarking and comparison of the cells grown ex vivo, and will assist with comparison of cells derived for clinical purposes. The methods that we describe here, and the resulting molecular classifier represent an important step towards addressing the more intractable questions of MSC identity, ontogenic relationships and function.

## Materials and Methods

Additional details are provided in the Extended Experimental Procedures

### Design of test and training datasets

We first carefully curated human MSC expression profiling experiments from GEO and ArrayExpress, evaluating each dataset to ensure samples met our criteria for primary data quality and a minimal MSC phenotype of CD105^+^, CD73^+^ and CD45^-^, with demonstrated differentiation to multiple mesenchymal lineages (see extended methods).

A review of public gene expression databases GEO and ArrayExpress identified 120 transcriptome datasets with MSC or stromal progenitors that also had accompanying publications. 35/120 datasets failed our QC criteria for data quality and were excluded from the study. Sixteen of the remaining 85 MSC datasets met our ‘gold standard’ criteria for accompanying phenotype and were included in our training set, together with 27 datasets containing cells from non-mesenchymal or non-stromal sources. Out of the 85 datasets, 37 included MSCs that lacked complete phenotype information and were assigned to a presumptive descriptor.

Non-MSC samples included immunophenotyped leukocyte subsets, primary and cultured differentiated cell types including fibroblasts from a wide range of sources, endothelial cells, pluripotent stem cells and cultured neurons. Our training set included 125 high-quality MSC and 510 non-MSC samples from 41 experiments, which had been profiled on 10 different microarray platforms (Supplementary Table S1 and S2). Finally, we used a combined dataset of 1291 samples, including 213 presumptive MSC (Supplementary Tables S6, S8, S9) to evaluate informativeness of the MSC signature on a variety of different cell types, extraction methods and growth conditions. All primary data are available from the Stemformatics stem cell resource. The underlying code for the statistical test is available as BootsPLS in the CRAN repository, and we have also made available the d3 code for the interactive MSC graph implemented in Stemformatics via the BioJS framework at http://biojs.io/d/biojs-vis-rohart-msc-test

### Identification of the 16-gene signature and assignation of a test sample to the MSC or non-MSC class

All microarray experiments analyzed in this paper were pre-processed using the R programming language (R Core Team, 2012). This involved a background correction of the raw data, a log_2_ and a YuGene transformation (*11*) which minimised batch effects caused by different microarray platforms (Supplementary Figure S4). The resulting data were then mapped to Ensembl gene ID to provide a common set of identifiers. A multivariate analysis, sparse Partial Least Square – Discriminant Analysis (Lê Cao et al., 2011), was used to select genes that discriminate MSC vs non-MSC on four components. The stability of each gene assigned to a component was assessed iteratively, using 200 randomised subsamplings of the training set in an internal 440-sample learning set and an internal 195-sample test set. We then determined the minimal informative signature by assessing the (i) stability and (ii) contribution to classification accuracy of all the genes selected at least twice over the 200 replications (Figure 1, Supplementary Figure S1). This reduced the signature to 16 key genes.

The final statistical model is fitted with the 16 signature genes on the training set and returns a score for each sample. The distribution of the training sample scores was used to determine an area of uncertainty [0.4337, 0.5169] (Supplementary Figure S1). By using 200 subsamplings of the training set, we recorded 200 scores for each sample, which enabled us to derive an individual 95% Confidence Interval (CI). A sample was assigned to the MSC class if the lower bound of its 95%CI is strictly higher than 0.5169. Similarly a non-MSC classification is given if the upper bound of the 95% CI was lower than 0.4337. Samples failing to meet these criteria were assigned as ‘unknown’.

### Network analysis

Twenty-six genes selected on component 1 equated to 18 proteins with a curated interaction in the Networkanalyst protein interaction database (which draws on the PPI database of the International Molecular Exchange (IMEx) consortium (*44*, *45*) These seed proteins were annotated to a shortest-path first-order network of 42 nodes and 52 PPI edges. Gene ontology analysis was assessed using hypergeometric mean against the Jan 2015 EBI UniProt GO library (*46*).

### Differential expression analysis

A linear mixed model with dataset as random effect was fitted for each gene for which both the mean of bone marrow samples and other sites were lower than the median of all gene expression values. This retains 16,903 genes. P-values were obtained by ANOVA and corrected for multiple testing with the Bonferroni-Hochberg procedure.

## Supplementary Files

**Table S1.** Datasets in the training set; accompanies Figure 1 and methods.

**Table S2**. Overview of the samples included in the training set, accompanies Figure 1 and methods.

**Table S3**. Datasets that contributed to the testing set: Accompanies Figure 2.

**Table S4**. Overview of the samples included in the test set with a presumptive call (MSC, non-MSC) and results of the Rohart test. Accompanies Figure 2.

**Table S5**. Overview of the samples included in the test set without a presumptive call (unknown) and results of the Rohart test. Accompanies Figure 2.

**Table S6**. Markers used in the common MSC immunophenotyping panel. Accompanies Figure 1 and Table 1.

**Table S7**. Members of the MSC signature network, seeded from component 1 genes and first-order interaction partners (BioGRID). Column 1 provides information on Gene Symbol (and ENSEMBL ID for seed members). Column 2 indicates Component or network membership and Probability of selection derived from sPLS-DA analysis. Disease annotations taken from the Mendelian Inheritance in Man (MIM) database. Accompanies Figure 2.

**Auxiliary Table S8**. Top Differentially expressed genes between MSC from bone marrow compared to other sites. Linear mixed model, FDR adjusted P<0.01

**Auxiliary Table S9**: Top Differentially expressed genes between MSC from different sites. Linear mixed model, FDR adjusted P<0.01.

**Supplemental Figure S1**: Overview of methods. Panel A, assessing stability of gene selection. Panel B, Determining a score between MSC and non-MSC.

**Supplemental Figure S2**: Box-whisker plots of the average expression of 32 common cell surface MSC markers across samples in the training dataset.

**Supplemental Figure S3**: Performance of the Rohart MSC signature across 3 time courses: A bone-marrow MSC adipogenesis; B bone-marrow MSC osteogenesis; C adipose-derived hASC reprogrammed to iPSC.

**Supplemental Figure S4**: Visualisation of training set clustering by PLS-DA plot to confirm lack of clustering driven by experimental batch. coloured by Stemformatics dataset ID (panel A); coloured by expression platform (Panel B); coloured by platform manufacturer (Panel C).

## References

1. A. J. Friedenstein, I. I. Piatetzky-Shapiro, K. V Petrakova, Osteogenesis in transplants of bone marrow cells. J. Embryol. Exp. Morphol.. 16, 381–390 (1966).

2. M. F. Pittenger et al., Multilineage potential of adult human mesenchymal stem cells. Science (80-.). 284, 143–147 (1999).

3. P. Bianco et al., The meaning, the sense and the significance: translating the science of mesenchymal stem cells into medicine. Nat Med. 19, 35–42 (2013).

4. D. G. Phinney, Functional heterogeneity of mesenchymal stem cells: Implications for cell therapy. J Cell Biochem. 113, 2806–2812 (2012).

5. A. Reinisch et al., Epigenetic and in vivo comparison of diverse MSC sources reveals an endochondral signature for human hematopoietic niche formation. Blood. 125, 249–260 (2014).

6. M. Crisan et al., A Perivascular Origin for Mesenchymal Stem Cells in Multiple Human Organs. Cell Stem Cell. 3, 301–313 (2008).

7. E. S. M. Lee, G. Bou-Gharios, E. Seppanen, K. Khosrotehrani, N. M. Fisk, Fetal stem cell microchimerism: natural-born healers or killers? Mol. Hum. Reprod.. 16, 869– 878 (2010).

8. R. M. Samsonraj et al., Establishing Criteria for Human Mesenchymal Stem Cell Potency. Stem Cells. 33, 1878–1891 (2015).

9. B. J. Sworder et al., Molecular profile of clonal strains of human skeletal stem/progenitor cells with different potencies. Stem Cell Res. 14, 297–306 (2015).

10. M. Dominici et al., Minimal criteria for defining multipotent mesenchymal stromal cells. The International Society for Cellular Therapy position statement. Cytotherapy. 8, 315–317 (2006).

11. K.-A. Lê Cao, F. Rohart, L. McHugh, O. Korn, C. A. Wells, YuGene: A simple approach to scale gene expression data derived from different platforms for integrated analyses. Genomics. 103, 239–251 (2014).

12. T. Çelik et al., PRRX1 is mutated in an otocephalic newborn infant conceived by consanguineous parents. Clin. Genet. 81 (2012), pp. 294–297.

13. E. Degenkolbe et al., A GDF5 Point Mutation Strikes Twice - Causing BDA1 and SYNS2. PLoS Genet. 9, e1003846 (2013).

14. B. R. Ali et al., Trafficking defects and loss of ligand binding are the underlying causes of all reported DDR2 missense mutations found in SMED-SL patients. Hum. Mol. Genet. 19, 2239–50 (2010).

15. I. Kawai et al., Discoidin domain receptor 2 (DDR2) regulates body size and fat metabolism in mice. Transgenic Res. 23, 165–175 (2014).

16. F. Zhang et al., Genome-wide copy number variation study and gene expression analysis identify ABI3BP as a susceptibility gene for Kashin–Beck disease. Hum. Genet. 133, 793–799 (2014).

17. N. Kaltz et al., Novel markers of mesenchymal stem cells defined by genome-wide gene expression analysis of stromal cells from different sources. Exp. Cell Res. 316, 2609–17 (2010).

18. R. J. Holley et al., Comparative Quantification of the Surfaceome of Human Multipotent Mesenchymal Progenitor Cells. Stem Cell Reports (2015), doi:10.1016/j.stemcr.2015.01.007.

19. Y. Koide et al., Two distinct stem cell lineages in murine bone marrow. Stem Cells. 25, 1213–1221 (2007).

20. F. M. Roversi et al., Serine protease inhibitor kunitz-type 2 is downregulated in myelodysplastic syndromes and modulates cell-cell adhesion. Stem Cells Dev. 23, 1109–20 (2014).

21. P. Charbord et al., A Systems Biology Approach for Defining the Molecular Framework of the Hematopoietic Stem Cell Niche. Cell Stem Cell. 15, 376–391 (2015).

22. S. Bae et al., Fibroblast activation protein alpha identifies mesenchymal stromal cells from human bone marrow. Br. J. Haematol. 142, 827–830 (2008).

23. S. Bae et al., Combined omics analysis identifies transmembrane 4 L6 family member 1 as a surface protein marker specific to human mesenchymal stem cells. Stem Cells Dev. 20, 197–203 (2011).

24. S. Gronthos, Molecular and cellular characterisation of highly purified stromal stem cells derived from human bone marrow. J. Cell Sci. 116 (2003), pp. 1827–1835.

25. H. M. Dansky et al., Adhesion of monocytes to arterial endothelium and initiation of atherosclerosis are critically dependent on vascular cell adhesion molecule-1 gene dosage. Arterioscler. Thromb. Vasc. Biol. 21, 1662–1667 (2001).

26. R. J. P. Skelton et al., SIRPA, VCAM1 and CD34 identify discrete lineages during early human cardiovascular development. Stem Cell Res. 13, 172–179 (2014).

27. H. Yoshida et al., KIAA1199, a deafness gene of unknown function, is a new hyaluronan binding protein involved in hyaluronan depolymerization. Proc. Natl. Acad. Sci.. 110, 5612–5617 (2013).

28. K. Zhang et al., The collagen receptor discoidin domain receptor 2 stabilizes SNAIL1 to facilitate breast cancer metastasis. Nat. Cell Biol. 15, 677–87 (2013).

29. F. Tremblay et al., Bidirectional modulation of adipogenesis by the secreted protein Ccdc80/DRO1/URB. J. Biol. Chem. 284, 8136–8147 (2009).

30. E. M. Walczak et al., Wnt-Signaling Inhibits Adrenal Steroidogenesis by Cell-Autonomous and Non-Cell-Autonomous Mechanisms. Mol. Endocrinol., me20141060 (2014).

31. B. Royer-Zemmour et al., Epileptic and developmental disorders of the speech cortex: Ligand/receptor interaction of wild-type and mutant SRPX2 with the plasminogen activator receptor uPAR. Hum. Mol. Genet. 17, 3617–3630 (2008).

32. D. Mrugala et al., Gene expression profile of multipotent mesenchymal stromal cells: Identification of pathways common to TGFbeta3/BMP2-induced chondrogenesis. Cloning Stem Cells. 11, 61–76 (2009).

33. F. S. Tedesco et al., Transplantation of genetically corrected human iPSC-derived progenitors in mice with limb-girdle muscular dystrophy. Sci. Transl. Med. 4, 140ra89 (2012).

34. M. N. Lim et al., Ex vivo expanded SSEA-4+ human limbal stromal cells are multipotent and do not express other embryonic stem cell markers. Mol. Vis. 18, 1289–300 (2012).

35. C. Wells et al., Stemformatics: Visualisation and sharing of stem cell gene expression. Stem Cell Res. 10, 387–395 (2012).

36. A. Papadimitropoulos et al., Expansion of human mesenchymal stromal cells from fresh bone marrow in a 3D scaffold-based system under direct perfusion. PLoS One. 9, e102359 (2014).

37. S. Suzuki et al., Properties and usefulness of aggregates of synovial mesenchymal stem cells as a source for cartilage regeneration. Arthritis Res. Ther. 14, R136 (2012).

38. S. Tanabe et al., Gene expression profiling of human mesenchymal stem cells for identification of novel markers in early- and late-stage cell culture. J. Biochem. 144, 399–408 (2008).

39. G. Walenda et al., TGF-beta1 does not induce senescence of multipotent mesenchymal stromal cells and has similar effects in early and late passages. PLoS One. 8, e77656 (2013).

40. J. M. Ryan et al., Transcriptional ontogeny of first trimester human fetal and placental mesenchymal stem cells: Gestational age versus niche. Genomics Data. 2, 382–385 (2014).

41. P. Benisch et al., The transcriptional profile of mesenchymal stem cell populations in primary osteoporosis is distinct and shows overexpression of osteogenic inhibitors. PLoS One. 7, e45142 (2012).

42. J. P. Martins et al., Towards an advanced therapy medicinal product based on mesenchymal stromal cells isolated from the umbilical cord tissue: quality and safety data. Stem Cell Res. Ther. 5, 9 (2014).

43. E. Seppanen et al., Distant mesenchymal progenitors contribute to skin wound healing and produce collagen: evidence from a murine fetal microchimerism model. PLoS One. 8, e62662 (2013).

44. S. Orchard et al., Protein interaction data curation: the International Molecular Exchange (IMEx) consortium. Nat Meth. 9, 345–350 (2012).

45. J. Xia, M. J. Benner, R. E. W. Hancock, NetworkAnalyst - integrative approaches for protein–protein interaction network analysis and visual exploration. Nucleic Acids Res.. 42, W167–W174 (2014).

46. R. P. Huntley et al., The GOA database: Gene Ontology annotation updates for 2015. Nucleic Acids Res.. 43, D1057–D1063 (2015).

